# Validation of the Tea Bag Index as a standard approach for assessing organic matter decomposition

**DOI:** 10.1101/2022.05.02.490129

**Authors:** Taiki Mori

## Abstract

The Tea Bag Index (TBI), a novel approach to assessing organic matter decomposition using commercial tea bags, has been increasingly utilized as a standard method in academic studies worldwide. This approach was designed to obtain an early-stage decomposition constant (*k*) indicative of early-stage decomposition rates and a litter stabilization factor (*S*) indicative of long-term carbon stability by using two types of teas—green and rooibos. However, despite the worldwide usage of the method, the accuracy of this approach has never been validated in terrestrial ecosystems. Here, the validity of this approach was tested by examining the two essential premises of the TBI using a laboratory incubation experiment. The first premise of the TBI—namely, that the unstabilized hydrolyzable fraction of green tea is mostly decomposed within 90 days—did not hold in the present study, which caused overestimations of the *S* of rooibos tea, as well as *k*. The second premise—namely, that the ratio of stabilized to total hydrolyzable fractions (i.e., *S*) of rooibos tea is equal to that of green tea—was also rejected, which resulted in substantial underestimations of the *S* of rooibos tea and *k*. Overall, the TBI largely underestimated the *S* of rooibos tea and *k* (more than 1.5 and 5 times smaller than those determined by time-series data, respectively). The present study suggests that time-series mass loss data of rooibos tea should be obtained to accurately determine *k*, rather than assuming that the *S* of rooibos tea is equal to that of green tea.

## Introduction

The Tea Bag Index (TBI) proposed by Keuskamp et al. (2013) is a novel method to assess organic matter decomposition using commercial tea bags as a standard material. This approach was designed to calculate a decomposition constant (*k*) and litter stabilization factor (*S*) using two types of teas, green and rooibos. The TBI was proposed to promote citizen science by providing a concise method to assess organic matter decomposition (Keuskamp et al., 2013), but the approach has been increasingly utilized as a standard method in the academic field worldwide (de Godoy Fernandes et al., 2021; Fung et al., 2022; Mueller et al., 2018; Petraglia et al., 2019; Pino et al., 2021).

In the TBI approach, *k* is defined as a decomposition rate constant of an asymptote model describing the decomposition curve of rooibos tea, which indicates early-stage decomposition rates. Meanwhile, *S* is defined as the ratio of stabilized to total hydrolyzable fractions of green tea, which was suggested to indicate long-term carbon stability (Fanin et al., 2020; Fujii et al., 2017). These two indexes can be calculated using a single pair of mass loss data of green and rooibos teas during a 90-day incubation (Fig. 1). This is possible because of the following two essential premises. First, the unstabilized hydrolyzable fraction of green tea is mostly decomposed within 90 days (note that the original paper suggested to check whether 90 days was sufficient, but 90 days are used without checking in most cases). In the framework of the TBI approach, the hydrolyzable fractions of green and rooibos teas (*H_g_* and *H_r_*, respectively, see Fig. 1) can be divided to the easily decomposable (*a_g_* and *a_r_*, respectively) and stabilized (i.e., recalcitrant) (*H_g_* – *a_g_* and *H_r_* – *a_r_*, respectively) fractions. Thus, the *S* of green and rooibos teas can be expressed as (*H_g_* – *a_g_*) / *H_g_* and (*H_r_* – *a_r_*) / *H_r_*, respectively. Under the first premise of the TBI approach (i.e., the unstabilized hydrolyzable fraction of green tea is mostly decomposed within 90 days), *S* can be determined using *H_g_* (0.842, determined by Keuskamp et al., 2013) and the mass loss ratio of green tea at the end of the 90-day incubation (*Mloss_G90*) as follows:

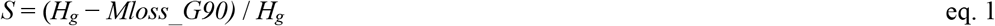

because under this premise, *a_g_* can be approximated to *Mloss_G90*. Notably, the TBI targets only the hydrolyzable fraction; thus, the decomposition of acid-insoluble fractions (1 – *H_g_* and 1 – *H_r_* for green and rooibos teas, respectively, see Fig. 1) is unable to be evaluated by this approach (Mori et al., 2021b). The second premise of the TBI is that the ratio of stabilized to total hydrolyzable fractions (i.e., *S*) of rooibos tea is equal to that of green tea. By using this premise, the stabilized hydrolyzable fraction of rooibos tea (*H_r_* – *a_r_*) can be determined by multiplying the total hydrolyzable fraction of rooibos tea (*H_r_*: 0.552, determined by Keuskamp et al., 2013) by the stabilization factor *S* determined with eq. 1. Thus, *a_r_* can be determined as follows:

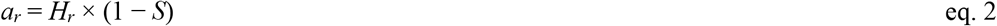

**Fig. 1.**
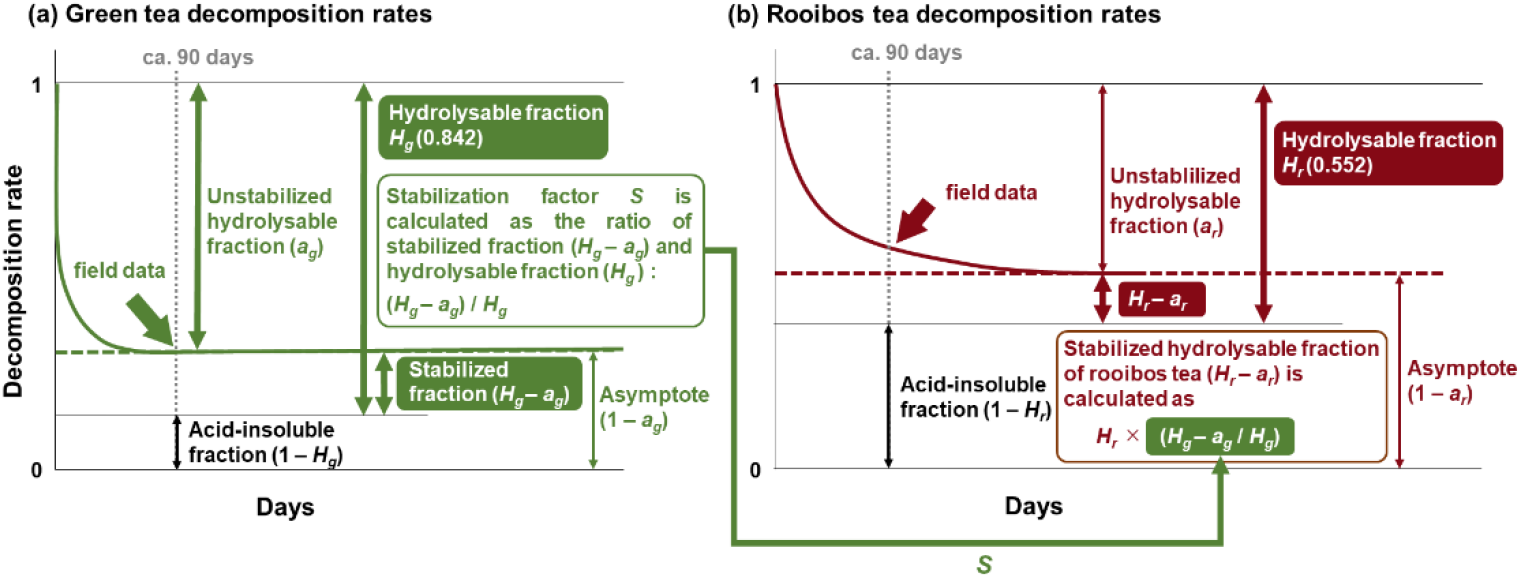
The Tea Bag Index (TBI) approach (modified from Mori et al., 2022). The decomposition curves of (a) green and (b) rooibos teas; *a_g_* and *a_r_* represent the unstabilized hydrolyzable fractions of green and rooibos tea, respectively. *H_g_* and *H_r_* represent the hydrolyzable fractions of green and rooibos tea, respectively.

The decomposition constant *k* can be determined by fitting an asymptote model to the mass loss ratio of rooibos tea at the 90th day (*Mloss_R90*) as follows (Fig. 1b):

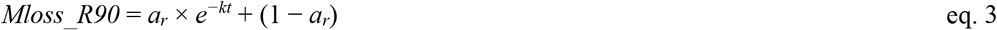

where *t* is the incubation period. Thus, the TBI approach provides a cost-effective and concise protocol to assess organic matter decomposition.

However, despite worldwide usage of the TBI, the accuracy of this approach has never been verified in terrestrial ecosystems. Recent studies have questioned the validity of this approach. The first premise of the TBI may not hold when green tea decomposition is slow (Keuskamp et al., 2013), which could overestimate both *k* and *S* (Mori et al., 2021b). Mori et al. (2022) demonstrated that *Mloss_R90* was larger than the decomposable fraction calculated using the first premise of the TBI and the green tea mass loss data (i.e., *a_r_* determined by the TBI) (see Fig. 1b), suggesting that the second premise is not always true. In aquatic ecosystems, the decomposition constant *k* calculated by the TBI was much larger than those determined by fitting time-series data of rooibos tea decomposition to an asymptote model (Mori, 2021; Mori et al., 2021c), possibly due to quick loss of the soluble fraction in the aquatic ecosystem. It is essential to test the validity of the TBI in terrestrial ecosystems. Teo et al. (2020) demonstrated that the decomposition curve of rooibos tea was within the range of TBI-based prediction, but their study was done under field conditions where environmental factors such as temperature and moisture are not constant.

In the present study, I performed a laboratory incubation experiment with controlled temperature and moisture levels to obtain time-series mass loss data of both green and rooibos teas. The validity of the TBI was tested by comparing the two indexes (hereafter *k_TBI* and *S_TBI*) and several key parameters (*a_g_* and *a_r_*; hereafter *a_g__TBI* and *a_r__TBI*) used in the approach with those determined by fitting an asymptote model to real data (hereafter *k_fitting, S_fitting_green, S_fitting_rooibos, a_g__fitting*, and *a_r__fitting*) as references [note that *S* can be determined using mass loss data for either green (*S_fitting_green*) or rooibos (*S_fitting_green*) tea]. The first premise of the TBI can be tested by comparing *a_g__TBI* with *a_g__ fitting*. The second premise can be tested by comparing *S_TBI* and *S_fitting_green* with *S_fitting_rooibos*. The incubation experiment was conducted under a low temperature condition to test the validity of the first premise under an unfavorable condition for tea decomposition.

## Materials and methods

### Tea bags

Following the TBI protocol (Keuskamp et al., 2013), Lipton green tea bags (EAN: 87 10,908 90359 5; Lipton) and Lipton rooibos tea bags (EAN: 87 22,700 18843 8; Lipton) were used. Since Lipton has changed the mesh from woven (0.25 mm) to nonwoven—but not the tea materials inside (Teatime 4 Science; http://www.teatime4science.org/)—I reproduced woven tea bags using 0.25-mm mesh and the tea materials from the current nonwoven tea bags (see Mori *et al*., 2021a).

### Incubation experiment

Surface soils (0–10 cm) were sampled from four randomly selected subplots established in a Japanese cedar [*Cryptomeria japonica* (L.f.) D.Don] plantation located at the Tatsudayama research site in Kumamoto, Japan (32.82 N, 130.73 E). Previous studies reported that the mean annual temperature and precipitation amounts at the research sites were 17.1°C and 1951 mm, respectively (Mori et al., 2021a) (data obtained from the Agro-Meteorological Grid Square Data of the National Agriculture and Food Research Organization). Sampled soils were sieved through a 4-mm sieve after the removal of large pieces of organic matter. A pair of tea bags (green and rooibos) was buried in 100 g of freshly sampled soil placed in a plastic (polyethylene terephthalate) bottle. The soil moisture was adjusted to 45% (w/w) because a previous study at the same site demonstrated that tea decomposition rates were highest around this moisture level (Mori et al., 2021a). Twenty bottles (4 subplots × 5 time-series) were placed in a cold room (3°C) for 5, 13, 23, 58, and 90 days. Water evaporation was prevented by covering bottles with polyethylene sheets (Mori et al., 2016). After incubation, tea bags were immediately oven-dried for few days to stop further decomposition. The dry weight of the tea bags was determined (oven-dried at 70°C for > 72 h) after the removal of surface soils and mesh.

### Statistics

Statistical analyses were performed using R version 4.1.1 (R Core Team, 2021). Parameters of the TBI approach (*k_TBI*, *S_TBI*, *a_g__TBI*, and *a_r__TBI*) were calculated following Keuskamp et al. (2013) (see also Fig. 1):

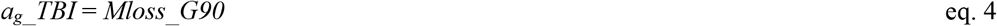

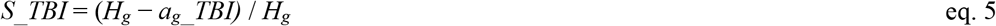

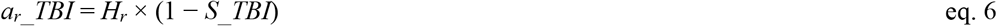

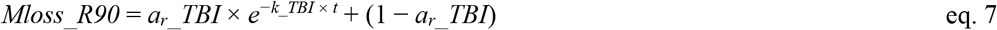

where *Mloss_G90* and *Mloss_R90* represent the mass loss ratios of green and rooibos teas at the end of the 90-day incubation, respectively, and *H_g_* (0.842) and *H_r_* (0.552) represent the hydrolyzable fractions of green tea and rooibos tea, respectively. Mass loss data for green and rooibos teas were fitted by asymptote models using nonlinear regression (“nls” library) to determine *k_fitting*, *a_g__fitting*, and *a_r__fitting* as follows:

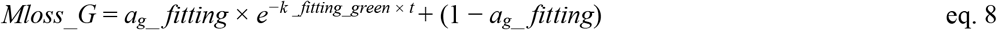

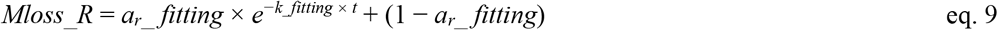

where *Mloss_G* and *Mloss_R* are the mass remaining ratios of green and rooibos teas, respectively, and *k_fitting_green* is the decomposition constant of green tea determined by fitting an asymptote model to time-series mass loss data for green tea. *S_fitting_geeen* and *S_fitting_rooibos* were determined as follows:

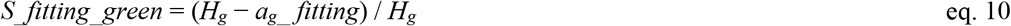

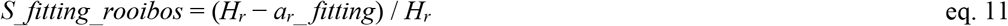

A linear mixed effect analysis was used to compare TBI-based parameters with parameters determined by fitting an asymptote model to real data.

## Results and discussion

The asymptote model fitted well to the time-series mass loss data of both green and rooibos teas (Figs. 2, 3). The fitting was much better than the single exponential model (Figs. S1, S2), justifying the usage of an asymptote model for predicting decomposition rates of teas in the TBI approach (Keuskamp et al., 2013).

**Fig. 2.**
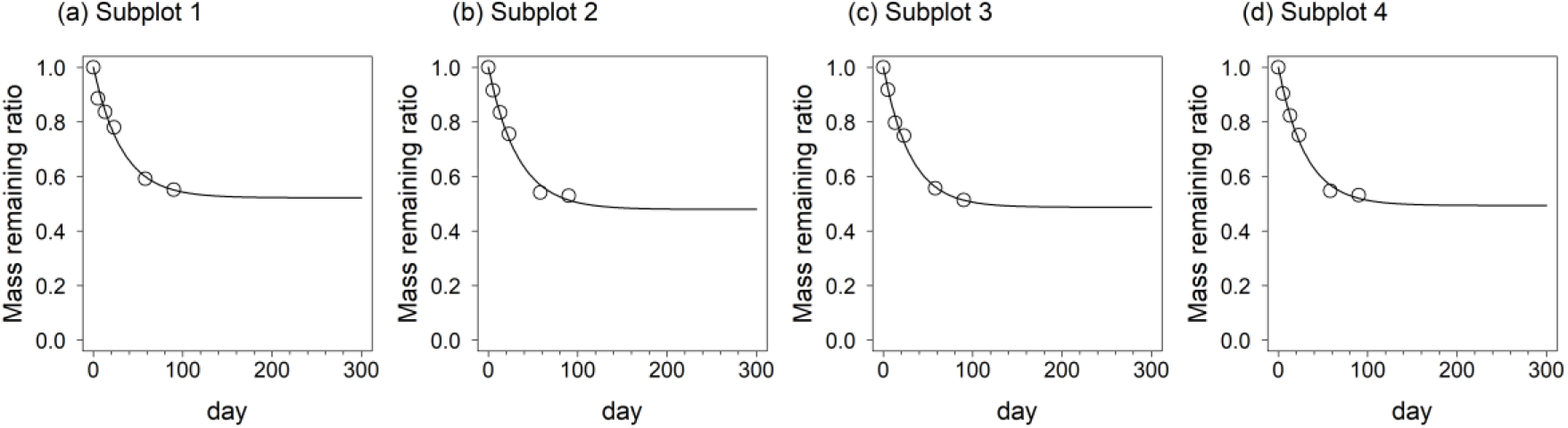
Relative mass of green tea remaining after a 90-day incubation. Open circles are mass loss ratios of green tea retrieved at 0, 5, 13, 23, 58, and 90 days after the start of incubation. Lines show fitting to the asymptote model: *W*(*t*) = *a* × *e^-kt^* + (1 – *a*), where *W*(*t*) is the mass remaining after incubation time *t*, *k* is the decomposition constant, and *a* is the unstabilized hydrolyzable fraction of green tea.

**Fig. 3.**
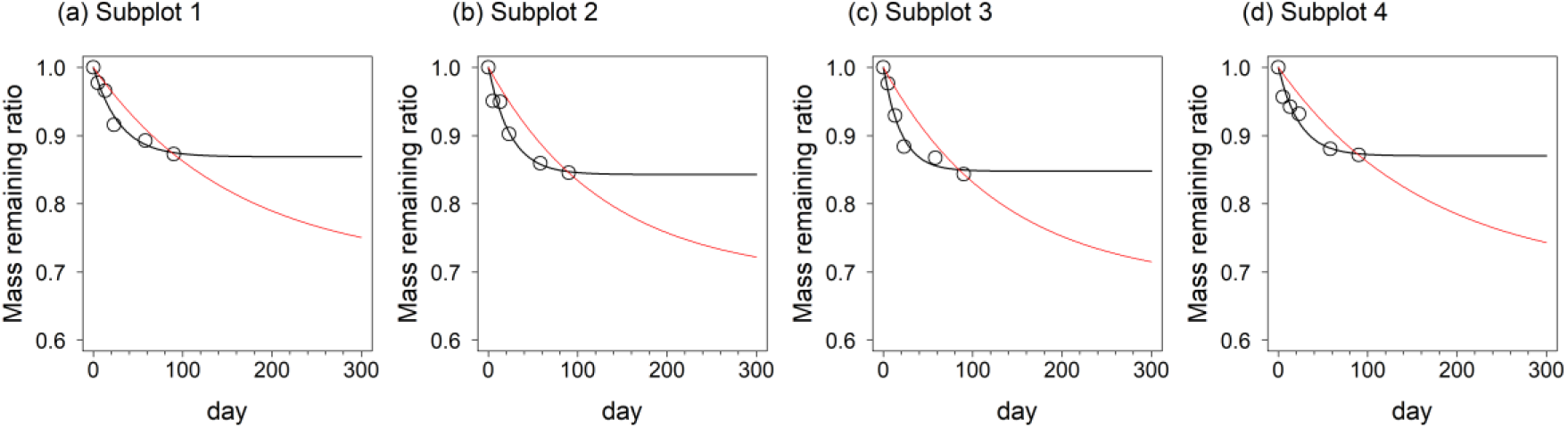
Relative mass of rooibos tea remaining after a 90-day incubation. Open circles are mass loss ratios of rooibos tea retrieved at 0, 5, 13, 23, 58, and 90 days after the start of incubation. Black lines show fitting to the asymptote model: *W*(*t*) = *a* × *e^-kt^* + (1 – *a*), where *W*(*t*) is the mass remaining after incubation time *t*, *k* is the decomposition constant, and *a* is the unstabilized hydrolyzable fraction of rooibos tea. Red lines show the asymptote model describing rooibos tea decomposition determined using the TBI.

In the present study, the first assumption of the TBI (i.e., the unstabilized hydrolyzable fraction of green tea [*a_g_*] can be approximated to the mass loss data of green tea at the end of the 90-day incubation) was not verified. Decomposition rates of green tea did not reach a plateau at the end of the 90-day incubation (Fig. 2), indicated by a slightly but significantly larger *a_g__TBI* (i.e., *a_g_* determined by the TBI approach) than *a_g__fitting* (i.e., *a_g_* determined by fitting) (Fig. 4). This was probably caused by the low temperature (3°C) used in this study, which could have retarded green tea decomposition. However, this overestimation of *a_g_* in the TBI approach was not large enough to influence the determination of *k*; *k_TBI* calculated using *a_g__fitting* was not significantly different from *k_TBI* calculated using *a_g__TBI* (Fig. 5c), although *a_r__TBI* and *S_TBI* were significantly larger and smaller, respectively, than those calculated using *a_g__fitting* (Fig. 5a, b).

**Fig. 4.**
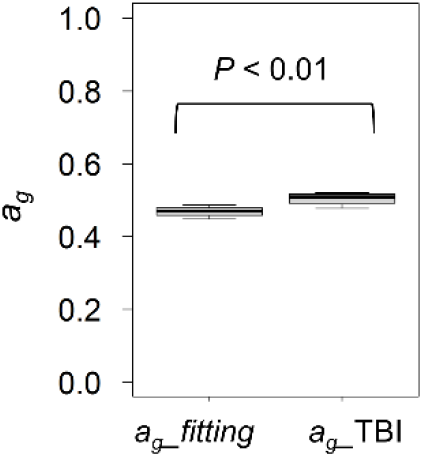
Comparison of the unstabilized hydrolyzable fraction of green tea (*a_g_*) determined by fitting an asymptote model to time-series mass loss data of green tea (*a_g__fitting*) and that determined by the TBI approach (*a_g__TBI*). A linear mixed effect analysis indicated *a_g__fitting* and *a_g__TBI* were significantly different (*P* < 0.01).

**Fig. 5.**
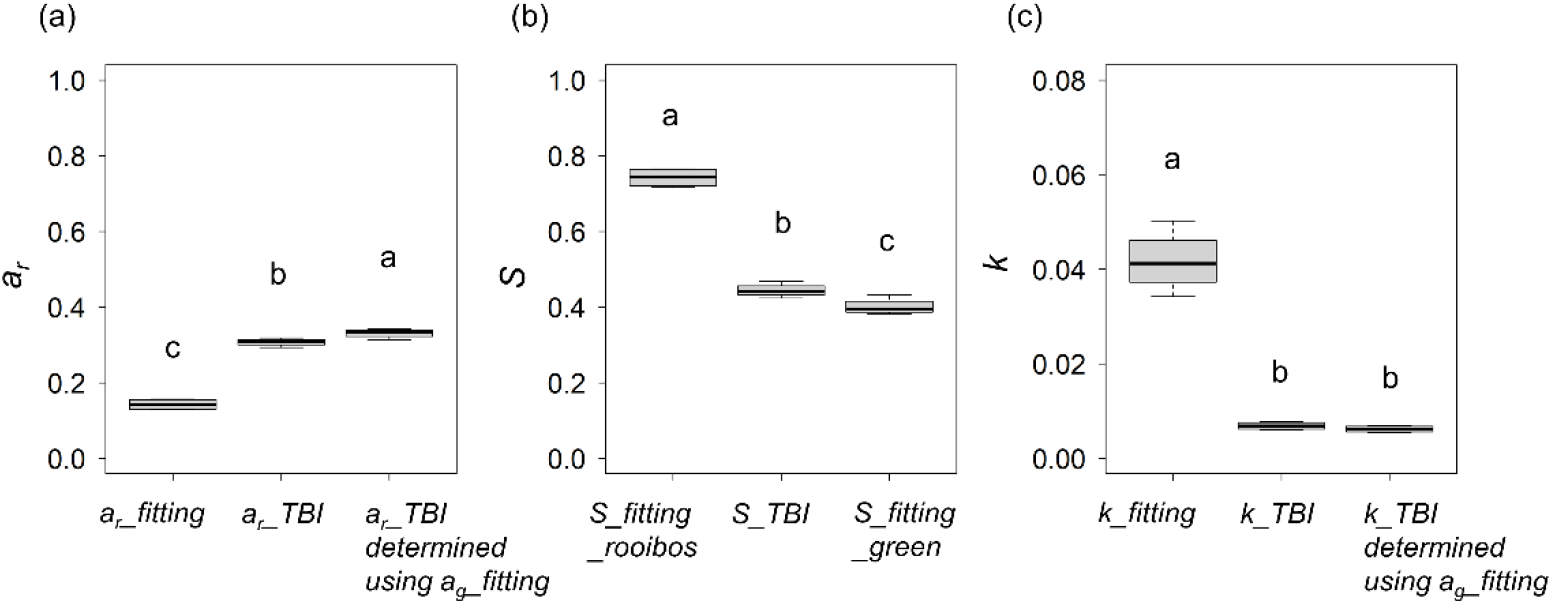
Comparison of the (a) easily decomposable fraction of rooibos tea (*a_r_*), (b) stabilization factor *S*, and (c) decomposition constant *k* determined by fitting an asymptote model to time-series mass loss data of green tea (*a_r__fitting, S_fitting_rooibos*, and *k_fitting*) to those determined by the TBI approach calculated using *a_g__TBI* as an unstabilized hydrolyzable fraction of green tea (*a_r__TBI*, *S_TBI*, and *k_TBI*), and those determined by the TBI approach calculated using *a_g__fitting* as an unstabilized hydrolyzable fraction of green tea (*a_r__TBI* determined using *a_g__fitting, S_fitting_green*, and *k_TBI* determined using *a_g__fitting*). Same letters in each figure indicate insignificant differences (*P* < 0.05, linear mixed effect analysis).

This study clearly shows that the second assumption of the TBI approach (i.e., the stabilization factor *S* of rooibos tea was equal to that of green tea) did not hold. The decomposable portion of the hydrolyzable fraction of rooibos tea (*a_r_*) determined by fitting an asymptote model to time-series mass loss data (*a_r__fitting*) was much larger than *a_r__TBI*, regardless of whether *a_r__TBI* was calculated using *a_g__TBI* or *a_g__fitting* (Fig. 5a). Accordingly, *S_fitting_rooibos* was also much larger than *S_TBI*, regardless of whether it was calculated using *a_g__TBI* or *a_g__fitting* (Fig. 5b). This result agrees with previous studies performed in aquatic ecosystems, which demonstrated that rooibos tea had a larger *S* than green tea (Mori, 2021, 2021c). Thus, the present study provides strong evidence for the assumptions in several previous studies: green tea and rooibos tea should have different *S* values because their chemical compositions are completely different (Mori et al., 2022, 2021b). The overestimated *ar* and underestimated *S* resulted in an underestimated *k* using the TBI approach (Fig. 5c). The *k_TBI* (0.0069 and 0.00622 when calculated using *ag_TBI* and *a_g__fitting*, respectively) was more than five times smaller than *k_fitting* (0.0417) (Fig. 5c). This substantially underestimated *k*, in combination with the underestimated *S*, caused large differences in the TBI-based decomposition curve compared to the fitting-based curve (Fig. 3).

This study is the first evaluation of the validity of the TBI for terrestrial ecosystems using a laboratory incubation experiment with controlled moisture and temperature conditions. Neither of the two essential premises of the TBI approach were verified in the incubation experiment, and the denial of the second premise (i.e., the stabilization factor *S* of rooibos tea is equal to that of green tea) is more important because the second premise should be independent of environmental conditions, while the first premise should be denied only under conditions unfavorable to decomposition (e.g., the low temperature condition in the present study). In addition, the denial of the second premise more strongly influenced the determination of *k* (Fig. 5c). In conclusion, I suggest that for academic studies, time-series decomposition data of rooibos tea should be obtained to accurately determine *k*, rather than assuming that the *S* of rooibos tea is equal to that of green tea, as suggested previously (Mori et al., 2022).

## Acknowledgments

I thank Ms Yumiko Sakamoto and Ms Akane Sakumori for their assistance on the laboratory work.

This research was funded by JSPS KAKENHI Grant Number JP19K15879.

## Supplemental materials

**Fig. S1.**
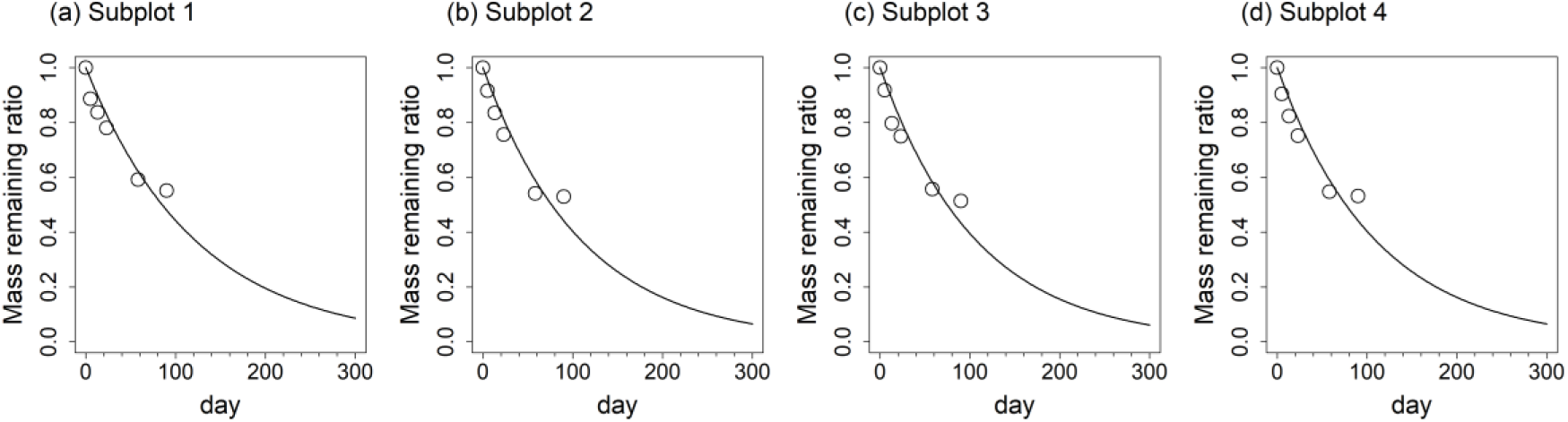
Relative mass of green tea remaining during 90-day incubation. Open circles are mass loss ratios of green tea retrieved at 0, 5, 13, 23, 58, and 90 days after the start of the incubation. Lines show fitting to asymptote model: *W*(*t*) = *e^-kt^*, where *W*(*t*) is the mass remaining after incubation time *t*, *k* is decomposition constant.

**Fig. S2.**
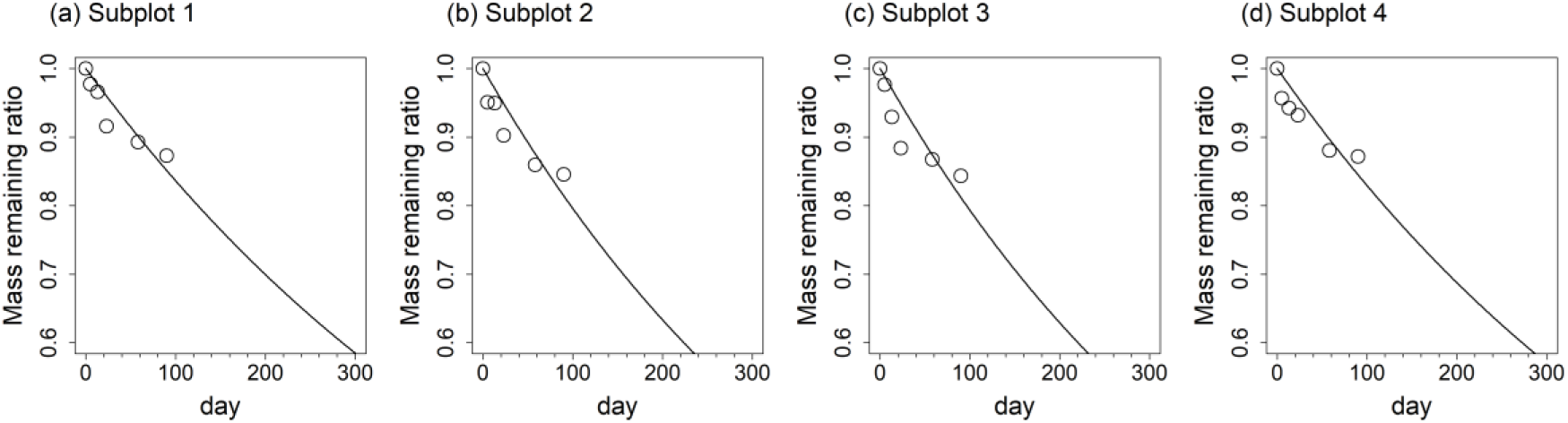
Relative mass of rooibos tea remaining during 90-day incubation. Open circles are mass loss ratios of rooibos tea retrieved at 0, 5, 13, 23, 58, and 90 days after the start of the incubation. Lines show fitting to asymptote model: *W*(*t*) = *e^-kt^*, where *W*(*t*) is the mass remaining after incubation time *t*, *k* is decomposition constant.

